# Understanding Performance Plateaus and Representation Collapse in 3D MRI Transformers Trained to Classify Brain Disease

**DOI:** 10.64898/2026.10.15.759510

**Authors:** Tamoghna Chattopadhyay, Kartik Shelar, Sophia I. Thomopoulos, Paul M. Thompson

## Abstract

In medical imaging AI, we commonly assume that diagnostic classifiers will increase in accuracy if they are trained on more data, but in reality, their performance often plateaus. To understand this behavior, we ran a spectral analysis of 3D vision transformers, a common supervised deep learning model that can be trained to classify disease based on 3D brain MRI. For simpler classification methods such as linear discriminant analysis, the classifier performance AUC(N), when trained on a sample of size N, is known to depend on the encoder’s eigenspectrum. Each class-discriminative direction in the latent space contributes only after the sample size N reaches a threshold N_i_ = λ_i_/α_i_^2^ - so the spectrum influences how performance scales with training data. Building on this theory, we show that supervised transformers compress almost all of their discriminative signal into one principal component (EVR@1 = 0.90–0.96): after this “representation collapse” happens, no further directions cross the threshold as training data increases, and accuracy plateaus. In principle, this plateau in performance could be due to either the encoder’s learned feature space or the difficulty of the task. To distinguish these cases, we trained three types of supervised 3D transformers (ViT3D, MINiT, NIT) and, as a non-collapsing alternative, a frozen self-supervised encoder (DINO), to classify people with dementia or mild cognitive impairment (MCI) versus cognitively normal (CN) individuals from 3D brain MRI, using data from the Alzheimer’s Disease Neuroimaging Initiative (ADNI). We froze each encoder and applied it to a new task, so that only the task difficulty varied. An “easy” task encoder (Dementia vs CN), transferred to the harder task (MCI vs CN), did not reproduce the plateau (MINiT threshold 9.2), but full retraining did (2758.5). The plateau is therefore due to the learned encoder, rather than task difficulty. Ranking the feature directions by the proportion of the disease signal they contain, we were able to identify a sign of representation collapse, i.e., a single dominant spike in the latent feature eigenspectrum. This spike was 11–26 times larger than predicted from the rest of the spectrum for the disease-trained (supervised) encoders, but was absent for DINO (0.66). To predict how transformer accuracy scales with the number of training scans, a large spike in the encoder spectrum suggests that the representation has collapsed and additional scans will not help; a small spike means that the signal is spread across many encoder latent space directions, and more training data should boost the classifier performance.

## 1. INTRODUCTION

A key practical question in neuroimaging AI is how much 3D MRI data we need to train a reliable brain disease classifier. Prior work on sample-size scaling laws in neuroimaging shows that prediction performance often plateaus as the amount of training data increases. Classical learning theory splits the classification error into an *irreducible component* (due to errors in the labeled data) and components that depend on the model itself and the amount of data available to train it [1-5]. For the simple case of linear discriminant analysis applied to two Gaussian-distributed classes in *p* dimensions, the recently proposed zeta law [9] explains how classification performance AUC(N) depends on the training sample size N. This theory is derived from inequalities that describe the finite-sample estimation error of the sample covariance, which needs to be estimated from the training data. The optimal discriminant depends on the inverse covariance matrix, but when we have few samples in a very high dimensional space (p ≫ N) this covariance cannot be estimated stably. From the sample Gram matrix, only the leading K(N) eigen-directions of the spectrum can be accurately resolved. K(N) is known as the *spectral horizon* for the sample size N [9]; the remaining covariance eigenvectors and eigenvalues are dominated by estimation error. In the linear case, classification accuracy at sample size N depends on a *truncated* Mahalanobis distance - the class-mean vector difference **d** projected onto these first K(N) reliable directions. A direction *i* enters this usable set only after its signal-to-noise ratio Nα_i_^2^/λ_i_ exceeds 1, that is, once N surpasses an admission threshold N_i_ = λ_i_/α_i_^2^, set jointly by the eigenvalue λ_i_ (the encoder’s covariance along direction *i*) and the signal projection α_i_ = u_i_^⊤^d (the alignment of **d** with that direction). As N grows, directions are admitted one at a time and AUC(N) rises gradually. If the leading threshold N_1_ already exceeds the largest feasible sample size, then no step is observable and classifier performance appears flat, i.e. it does not get better with more training data.

Applying this theoretical model to neuroimaging data, in [6] we trained supervised vision transformer (ViT) encoders on 3D brain MRI scans from ADNI [7] to classify dementia cases versus cognitively normal (CN) individuals. The deep learning models, during training, tended to collapse their CLS embedding spaces to near-rank-1 geometry. A single principal component in the final embedding space captures 90–96% of the total variance in the data, and the dominant admission threshold N_1_* falls below 1. Because of this collapse to a single rank-1 component, the classifier performance AUC(N) remained flat across the training range N ∈ [50, 4000]. Using a DINO ViT-B/16 encoder (adapted to handle 3D images) as a positive control [8], we showed in [6] that collapse depends on the training objective rather than the transformer architecture: the same transformer backbone collapses under a supervised objective but not under DINO’s self-supervised one, so it is the objective, not the use of a transformer, that drives the collapse. The present paper examines this further: by freezing a trained encoder and swapping only the contrast, we separate the objective’s effect from the effect of the architecture and from task difficulty.

This leads to a practical question: *if the training objective drives the collapse, does the difficulty of the task also influence it?* If a harder task simply raises the leading admission threshold N_1_ beyond the current sample size, more scans will boost performance eventually; but if the collapse is a fixed property of the learned representation, independent of the task, additional data will not help after a certain point. Even so, retraining a neural network alters two factors at once - the encoder’s covariance λ and the signal projection α of the class-mean difference onto each eigendirection. To distinguish these two causes, we freeze a trained encoder and reapply it to a new contrast, holding the encoder spectrum fixed by construction so that only the disease loading α can vary in the embedding. Using this frozen-encoder design, we test whether the estimated thresholds carry real signal or lie within their permutation null. We use these tests to identify the cause of the representation collapse to the transformer training rather than an attention pooling step that is often used in transformer design. We test these effects in experiments with three supervised transformer models (ViT3D, MINiT, NIT) and a self-supervised DINO architecture on the ADNI dataset (CN/Dementia and MCI/CN). We study which factors influence the scaling behavior: the training objective, task difficulty, or the learned representation that the two produce together. DINO is trained without using disease labels (self-supervised), so by using it as a control, we show that the collapse comes from the supervised training objective, not from the transformer architecture itself.

This study has three main contributions. First, we study whether collecting more scans will improve a brain MRI classifier and show that this does not only depend on the task; it is a property of the representation the model has learned. Using a “frozen” encoder that fixes the feature space while the task varies, we show that the scaling behavior depends on the trained encoder, not on the difficulty of the classification. As a result, two deep learning models trained on the same task can have different data scaling laws. Second, we show how to use the shape of the feature spectrum to determine whether accuracy will increase with more training data. A key sign of a performance plateau is a representation collapse. We show how to identify this as a single dominant rank-1 spike in the data encoder covariance spectrum. We find that this single dominant rank-1 spike is large for the supervised disease-trained encoders but is not found for a self-supervised encoder, DINO. Third, this yields a practical planning approach when we are training diagnostic classifiers for brain MRI or even for other types of medical data. Measured on a small pilot sample, a rank-1 spike in the encoder spectrum indicates whether a trained model is in an active scaling regime (where more data will help), or a saturated one (where it will not). Our spectral model of learning curves offers an alternative to trial-and-error inspection of the AUC(N) scaling curve. Spectral analysis of the encoder provides a helpful test that we can perform when training a diagnostic classifier, and when waiting for new data to come in. By identifying a rank-1 spike in a pilot sample, we can help to identify whether the (architecture, task) pair is in an active or saturated scaling regime.

## 2. BACKGROUND

### A. The Linear Classifier setting and the Zeta Law

The recently proposed *zeta law* framework [9] analyzes a linear two-class (binary) classification problem in high dimensions. A classifier trained on N labeled examples in p≫N dimensions cannot stably estimate the full covariance of the encoded data. Only the leading K(N) eigen-directions of the sample covariance Σ^=X^⊤^X/(N−1) can be accurately resolved above sampling noise. When p≫N, we can estimate these directions efficiently through the corresponding N×N

Gram matrix XX^⊤^. Classification therefore uses only the projection of the class-mean vector difference **d** onto those K(N) directions - a *truncated Mahalanobis distance*. Direction *i* becomes usable once its signal-to-noise ratio Nα_i_^2^/λ_i_ exceeds 1, i.e. once N > λ_i_/α_i_^2^. We call N_i_ = λ_i_/α_i_^2^ the *admission threshold* of direction i: the sample size at which that direction begins to contribute a net positive signal. Directions are admitted one at a time as N grows, so the expected value of AUC(N) across population samples of size N is predicted to rise gradually. If N_1_ falls below the smallest feasible sample size, the entire staircase (of performance gains with N) is invisible and performance appears flat.

Formally, let Σ = UΛU^⊤^ be the PCA covariance of the data embeddings, with eigenvalues λ_i_ ∝ i^−β^ (spectral slope β, estimated by OLS over i = 1–50). Σ = UΛU^⊤^ is essentially PCA written with matrices. U holds the principal directions (eigenvectors u_i_) and Λ holds their variances (eigenvalues λ_i_). Sorted, the λ_i_ fall off like i^−β^, and the exponent β (the ‘spectral slope’) says how fast — bigger β means variance is packed into fewer directions. d is the vector from the mean healthy embedding to the mean diseased embedding: the direction that separates the classes. α_i_ = u_i_^⊤^d measures how much of that separating direction lies along PCA direction i, and these loadings also fade, like i^−^η (exponent η). We let α_i_ = u_i_^⊤^d, with α_i_ ∝ i^−^η (signal-decay exponent η). For a linear classifier, performance follows AUC(N) = Φ(d□(k(N))/√2), where d□^2^(k) = Σ_i_≤1 α_i_^2^/λ_i_ is the truncated Mahalanobis distance and k(N) = max{i : Nα_i_^2^/λ_i_ > 1}. This formula holds under an equal-covariance Gaussian model for the optimal LDA score restricted to the retained subspace. Φ is the normal-curve function that converts a separation distance into an AUC. d□^2^(k) adds up the usable signal-to-noise α_i_^2^/λ_i_ over the first k directions (the ‘truncated Mahalanobis distance’), and k(N) counts how many directions are switched on at sample size N. When 2η > β, admission is ordered and directions enter in increasing index, producing a regular staircase. So if 2η > β (the squared signal fades faster than the variance), directions switch on neatly in order 1, 2, 3, …, giving the clean ‘staircase.’ Throughout, α_i_ denotes the signal projection and ρ denotes the L(k) power-law exponent, L(k) = A/k^ρ + δ. The spectral slope should be described as the negative slope from a regression of log λ_i_ on log i.

### B. Spectral Collapse in Supervised Encoders

Although the spectral analysis of performance scaling depends on arguments from the linear case, in [6] we began testing whether these predictions also hold for transformers, which are nonlinear. We expect them to be applicable as heuristics, as the nonlinearity is only in the encoder, which maps inputs to embeddings. Once the embeddings are frozen, class separability depends only on the geometry of the vector embedding space. For a linear classifier in this embedding space, this separability still depends on the class-mean vector difference **d** and the covariance Σ, i.e., it depends on the projections α_i_ = u_i_^⊤^d and the eigenvalues λ_i_. The encoder’s nonlinearity sets the values of β and η, but for fixed β and η the linear staircase still describes how AUC grows with N. The baseline of our prior paper [6] is used throughout. Recall that β is the spectral slope: how fast the embedding eigenvalues λ_i_ decay, with larger β meaning the variance is concentrated in fewer directions. The effective rank counts how many directions carry meaningful variance (a value near 1 means almost all variance sits in a single direction), and EVR@1 is the fraction of total variance explained by the first principal component alone. N_1_* is the sample size at which the first direction becomes detectable, i.e., the smallest N for which Nα_1_^2^/λ_1_ > 1; if N_1_* is below any realistic training set, the first step of the staircase cannot be seen and AUC(N) is flat.

In our prior work [6], for Dementia vs CN classification on ADNI, we found that ViT3D, MINiT and NIT showed a spectral slope β = 2.00–2.31, effective rank 1.27–1.72, EVR@1 = 0.90–0.96, and N_1_* = 0.3–0.5. As these N_1_* values fall below any feasible training set, the AUC(N) staircase was not observable. The DINO positive control instead achieved β = 1.148 (a slower spectral decay), effective rank ≈ 52, and a rising AUC(N) curve, showing that the scaling law depends on the type of encoder used. Together these results establish that the near-rank-1 collapse is present for the supervised encoders but not found for the self-supervised one; the present work identifies which factors - the objective, contrast, or representation - are responsible for this behavior.

### C. Neural collapse and contrast separability

Neural scaling laws characterize performance as power laws of the training dataset size (or of computing scale) [13,14]. Spectral analysis of representation geometry has been applied to study neural collapse [15,16] and feature isotropy [12,17] in self-supervised learning. ‘Feature isotropy’ means that variance spread evenly across directions in the embedding space of the data encoder (the opposite of collapse). In neuroimaging, sample-size requirements are often estimated via Cohen’s *d* on brain summary statistics [2]; spectral power analysis offers a complementary approach extended to higher-dimensional feature sets. The MCI vs CN discrimination task is well-known to be harder than classifying Dementia vs. CN, with smaller expected effect sizes and near-chance performance for classifiers based solely on structural MRI [18]. ‘Neural collapse’ is a known end-of-training phenomenon where supervised training pulls each class’s features onto its class average and arranges the class centers into a tidy geometric pattern (a ‘simplex’). [19]. For two classes this predicts exactly one dominant direction - a rank-1 spike sitting above the otherwise smooth eigenvalue trend, not a change in the slope of the trend. A key corollary is that collapse should occur only when the training contrast is linearly fittable. A separable contrast can be collapsed onto its between-class direction, whereas a non-separable contrast cannot.

## 3. METHODS

### A. Dataset and Splits

We analyzed publicly available data from the Alzheimer’s Disease Neuroimaging Initiative (ADNI) [7], selecting 3D T1-weighted volumetric brain MRI data from participants who were cognitively normal (CN) or had mild cognitive impairment (MCI). For completeness, we also report Dementia vs. Control data from our prior study [6]. Participants were split 70/15/15 at the subject level, stratified by diagnosis and sex, yielding 4,057 training scans, 877 validation scans, and 858 test scans. MCI was the majority class (3,479 vs 2,313 CN scans) with a positive class weight = 0.6785. Details are in **Table 1**. All spectral quantities - eigenbasis, class-mean difference **d**, α_i_ and λ_i_ - were computed on the training split (N = 4,057) from the CLS token taken before the classification head; as ‘plug-in’ Mahalanobis distance inflation scales with 1/n. In this context, ‘plug-in Mahalanobis inflation’ means: if we estimate the class separation from finite samples (‘plug in’ the sample estimates), the Mahalanobis distance is slightly biased upward, because random noise in a finite sample always adds a little apparent separation. That inflation shrinks in proportion to 1/n, and the more samples we compute the distance from, the smaller the bias. As the size of the bias depends on how many samples were used, we fix the extraction set (the training split, N = 4,057), so the bias is identical everywhere and comparisons are fair.

**Table 1.** Data Split for MCI vs CN.

| **Table 1.** Data Split for MCI vs CN. |  |  |  |  |  |
| --- | --- | --- | --- | --- | --- |
| **Split** | **Scans** | **Subjects** | **CN** | **MCI** | **Age** |
| Train | 4,057 | 959 | 1,640 | 2,417 | $74.7 \pm 7.1$ |
| Validation | 877 | 206 | 330 | 547 | $74.3 \pm 7.3$ |
| Test | 858 | 206 | 343 | 515 | $75.6 \pm 7.4$ |
| Total | 5,792 | 1,371 | 2,313 | 3,479 | $74.7 \pm 7.2$ |

### B. Architectures and Training

Three supervised architectures from [6] were evaluated: ViT3D (5.85M params, 6 layers, 8 heads, D = 256), MINiT (3.27M, block-wise encoder with ABMIL aggregation, D = 256), and NIT (5.85M, D = 256). All receive 96×112×96 T1-MRI volumes tokenized into 252 patches of 16^3^ voxels. Training hyperparameters were identical to [6]: AdamW, lr = 1×10^−4^ with cosine annealing and 5-epoch warmup, effective batch = 16, max 100 epochs, early stopping patience = 15 on val AUROC, class-weighted BCE (pos_weight = 0.6785 for MCI/CN). The DINO ViT-B/16 encoder with 3D-inflated patch embeddings and frozen backbone, described fully in [6], appears as a validation input for the zeta law forward model and is not re-trained here. For the seed-variance and frozen-encoder analyses, we trained 10 seeds per architecture per task (54 runs) and stored their CLS embeddings.

### C. Spectral Pipeline and Estimators

Our spectral analysis pipeline from [6] provides the fundamental measurements for fitting eigenvalue decay laws (OLS fit of log λ_k_ versus log *k* over the range *k* = 1–50 for β, effective rank and EVR@k; the Mahalanobis spectrum α_i_ = u_i_^⊤^d; admission thresholds N_i_ = λ_i_/α_i_^2^; the large-sample prediction AUC_pred = Φ(d□/√2); the power-law fit L(k) = A/k^ρ^ + δ; and the k(N) subsampling sweep). A GPU implementation of SVD was used to replace the CPU TruncatedSVD of [6] for the larger dataset. For our identification strategy, we used five estimators:

a. **Permutation null**. CN/MCI labels were shuffled within the embedding set 500 times, preserving class sizes; d, α_i_, d□ and N_1_ were recomputed on the fixed (label-independent) eigenbasis, giving the distribution of N_1_ under the setting of “no signal, same covariance.”
b. **Seed variance and split-half stability**. Spectral quantities were recomputed for each of the 10 seeds, and over 100 partitions the eigenbasis is estimated on one subject half, and **d** on a disjoint half.
c. **Cross-fitted debiasing**. As 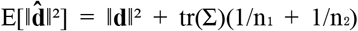 \, plug-in d□ is upward-biased. We compute three estimators side by side: plug-in; a subject-level 3-way cross-fit (basis, 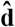 and evaluation on disjoint folds); and the closed-form correction 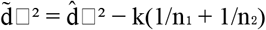 .
d. **Frozen-encoder 2×2**. A full encoder × contrast design (below) with subject-level leakage control, run in both a frozen-AD basis and a native basis, with principal angles computed between them. By subject-level leakage control, we mean that when a frozen encoder is applied to the other cohort (cells 2 and 4), any subject whose scans were used to train that encoder should be excluded from the evaluation set, so the transferred α is measured only on unseen subjects and cannot reflect memorization. Exclusion is at the subject level rather than the scan level because a second scan of the same subject shares the same anatomy.
e. **Spike ratio**. A bulk slope β_bulk is fit over i = 2–50 (excluding PC1); the spike ratio 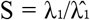 (bulk) is computed with a 200× subject-level bootstrap, alongside the analytic check EVR_1_ = 1/ζ_D(β), where ζ_D(β) = Σ_{i=1}^{D} i^−β^ is the Riemann zeta function truncated to the D embedding dimensions (D = 256 for the supervised encoders, 768 for DINO); this is the PC1 variance share that a pure power law of slope β would produce, so a large gap between observed EVR@1 and 1/ζ_D(β) is the same spike measured from the variance side. Here EVR_1_ = 1/ζ_D(β) is a theoretical cross-check, as a perfectly smooth power-law spectrum predicts a specific fraction of the variance in PC1, and we obtain the Riemann zeta function ζ when we sum the power law.

## 4. RESULTS

### 4.1 Spectral geometry across training objective and contrast

Table 2 summarizes the spectral parameters for both contrasts and all encoders. The between-condition differences depend on the covariance spectrum λ: the spectral slope β, effective rank and EVR@1 all change between CN/Dementia and MCI/CN and between the supervised encoders and DINO. On the more difficult MCI contrast, the geometry does not de-collapse uniformly as MINiT remains near-rank-1 (EVR@1 = 0.94, effective rank ≈ 1.45) while ViT3D and NIT partially spread (0.77, 0.65), so the collapse does not depend only on task difficulty. As a model retrained on a new contrast changes both its covariance λ and its signal projection α, a comparison across retrained models cannot attribute a change to either factor alone; to isolate the main causes, we hold one factor fixed, using a frozen-encoder design. Application of the pipeline to MCI vs CN also yields, for MINiT, an apparent admission threshold N_1_ = λ_1_/α_1_^2^ = 283.6 that should place a staircase step within an observable sample-size range. As N_1_* = λ_1_/α_1_^2^ has the small quantity α_1_^2^ in its denominator, we find through bootstrapping that this parameter is large and unstable when the signal projection α_1_ is near zero. Because it is a ratio, it is also unchanged if the whole embedding is rescaled (λ → c^2^λ, α → cα), so a single N_1_* value carries no information about the absolute size of the covariance or the signal. We therefore examine the reliability of N_1_* and express it in terms of its underlying components (tr Σ, λ_1_, ║d║^2^, α_1_) in **Table 3** so that a given estimate of N_1_* can be assessed based on whether α_1_ has collapsed or the overall scale has shrunk.

### 4.2 Frozen-encoder experiments: the spectral geometry depends on the encoder, not the classification contrast

To understand whether the spectral geometry depends *only* on the type of encoder or also on the task contrast (the classes that need to be separated), we held the encoder fixed and swapped only the classification contrast. In this case, λ_k is identical by construction and only α_k (the classifier’s loading of discriminative energy onto the encoder modes) can be altered. In this 2×2 experimental design, we treat N_1_* as an ordinal, distributional quantity; comparing seed ranges rather than point values; and we check every observation about N_1_* against the scale-free geometry (EVR@1, spike ratio), for reasons of identifiability noted in Section 4.5. We write E_AD for the encoder trained on the Dementia-vs-CN contrast and E_MCI for the same architecture trained on the MCI-vs-CN contrast; each is then evaluated on both contrasts. Four cells were run per architecture: (1) the E_AD encoder on Dem−CN (baseline); (2) E_AD encoder on MCI−CN (α-only, λ frozen); (3) the E_MCI encoder on MCI−CN (the retrained condition); and (4) E_MCI encoder on Dem−CN (reverse transfer). Cells 2 and 4 are forward passes of a frozen encoder over the other cohort. Subject-level leakage control excludes any subject in the encoder’s training split. To confirm the two covariances share a common leading geometry, we compute the principal (canonical) angles between their top-k eigen-subspaces: writing U_k and V_k for the matrices of top-k eigenvectors (orthonormal columns) of the frozen-AD and native MCI-cohort covariances, the principal angles are θ_j = arccos(σ_j), where σ_j are the singular values of U_k^T^ V_k [9]. The smallest angle θ_1_ is the closest alignment between the subspaces and the largest θ_k the worst. The leading eigenvectors align to within θ_1_ = 0.1–0.3° (k = 1), and for the top-five subspace (k = 5) the largest principal angle is θ_5_ ≲ 5°, confirming that cell 2 is a genuine α-only measurement rather than a cohort-covariance artifact. **Table 3**. gives the full absolute-scale decomposition (seed 42, cross-fit, native basis; cells 2 and 4 leakage-controlled), and **Figure 1** shows the resulting thresholds. The most important comparison is cell 2 vs cell 3. Because N_1_* is not a stable point estimate in this regime, we base the comparison on the whole seed distribution and on the scale-free geometry rather than on single values. Across all 10 seeds the MINiT cell-2 and cell-3 distributions do not overlap: cell 2 spans 5.4–17.8 while cell 3 spans 28–22,949 (representative seed-42 values 9.2 and 2758.5). Routing the hard contrast through the frozen AD encoder therefore does not reproduce the elevated threshold produced by retraining. The same separation appears in quantities that are stable and estimator-independent. EVR@1 remains ≈0.94 across cells 1–4 while α_1_ and the overall scale change, and the rank-1 spike remains large in cell 2 (S ≈ 10) - so the geometry that follows the encoder is visible without relying on N_1_*. Because only α can differ between cells 2 and 3, the effect is not carried by α alone. The reverse transfer confirms it: the MCI-trained encoder on the easy contrast still gives N_1_ = 39.8 (cell 4) versus 0.36 for the same easy contrast on the AD encoder (cell 1). The encoder, not the contrast, carries the regime, and across all 10 seeds the MINiT cell-2 ≪ cell-3 ordering holds without exception (cell 2 spans 5.4–17.8, never approaching cell 3’s 28–22,949).

**Table 2.**
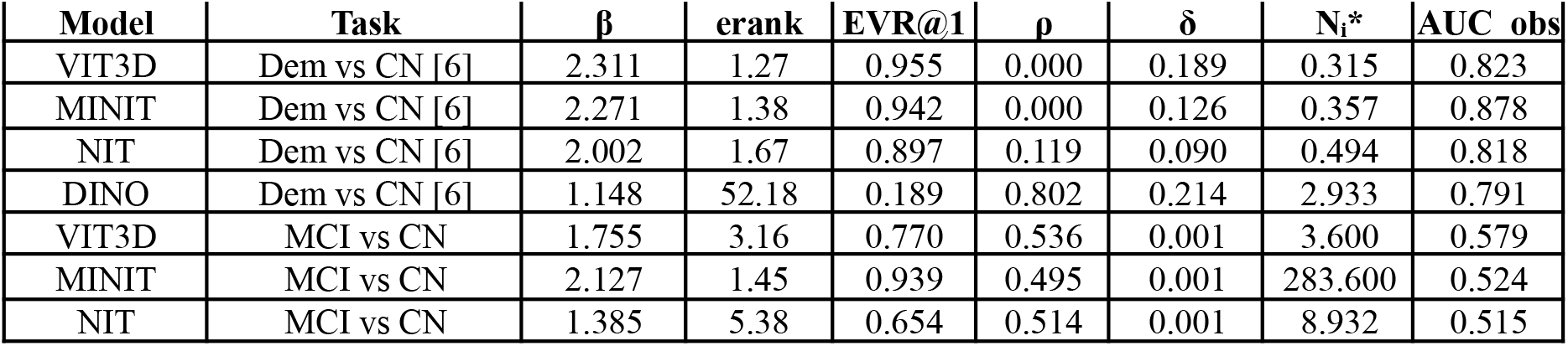
Spectral parameters (plug-in estimator, training split). β: spectral slope. ρ: L(k) power-law exponent. δ: error floor. N_1_*: PC1 admission threshold. CN vs Dementia and DINO baseline values follow [6]. DINO effective rank/EVR@1 use the full-D (768) convention

**Table 3.** Frozen-encoder 2×2 decomposition (seed 42, cross-fit, native basis; cells 2 and 4 leakage-controlled). Cell key: 1 = E_AD·Dem−CN, 2 = E_AD·MCI−CN, 3 = E_MCI·MCI−CN, 4 = E_MCI·Dem−CN.

| Model | Cell | $\text{tr}\Sigma$ | $\lambda_1$ | $EVR@1$ | $\ d\ ^2$ | $\alpha_1$ | $N_1^*$ | $d_\square$ |
| --- | --- | --- | --- | --- | --- | --- | --- | --- |
| MINiT | 1: $E\_AD$-Dem-CN | 17.06 | 16.06 | 0.942 | 45.05 | 6.710 | 0.357 | 1.936 |
| MINiT | 2: $E\_AD$-MCI-CN | 11.20 | 10.22 | 0.913 | 1.326 | 1.124 | 9.191 | 0.951 |
| MINiT | 3: $E\_MCI$-MCI-CN | 0.049 | 0.046 | 0.939 | 0.0003 | 0.0141 | 2758.5 | 0.635 |
| MINiT | 4: $E\_MCI$-Dem-CN | 0.047 | 0.044 | 0.935 | 0.0022 | 0.0412 | 39.84 | 1.208 |
| ViT3D | 1: $E\_AD$-Dem-CN | 94.58 | 90.33 | 0.955 | 286.98 | 16.93 | 0.316 | 2.097 |
| ViT3D | 2: $E\_AD$-MCI-CN | 58.81 | 55.02 | 0.936 | 1.823 | 1.228 | 89.19 | 0.909 |
| ViT3D | 3: $E\_MCI$-MCI-CN | 0.371 | 0.283 | 0.769 | 0.0805 | 0.2804 | 3.726 | 0.994 |
| ViT3D | 4: $E\_MCI$-Dem-CN | 0.393 | 0.304 | 0.777 | 0.1521 | 0.3835 | 2.119 | 1.404 |
| NIT | 1: $E\_AD$-Dem-CN | 21.57 | 19.36 | 0.899 | 39.35 | 6.262 | 0.497 | 1.787 |
| NIT | 2: $E\_AD$-MCI-CN | 14.50 | 12.42 | 0.858 | 1.485 | 1.193 | 9.491 | 0.920 |
| NIT | 3: $E\_MCI$-MCI-CN | 0.063 | 0.040 | 0.649 | 0.0054 | 0.0667 | 9.524 | 1.050 |
| NIT | 4: $E\_MCI$-Dem-CN | 0.078 | 0.055 | 0.715 | 0.0092 | 0.0854 | 8.953 | 1.481 |

**Figure 1.**
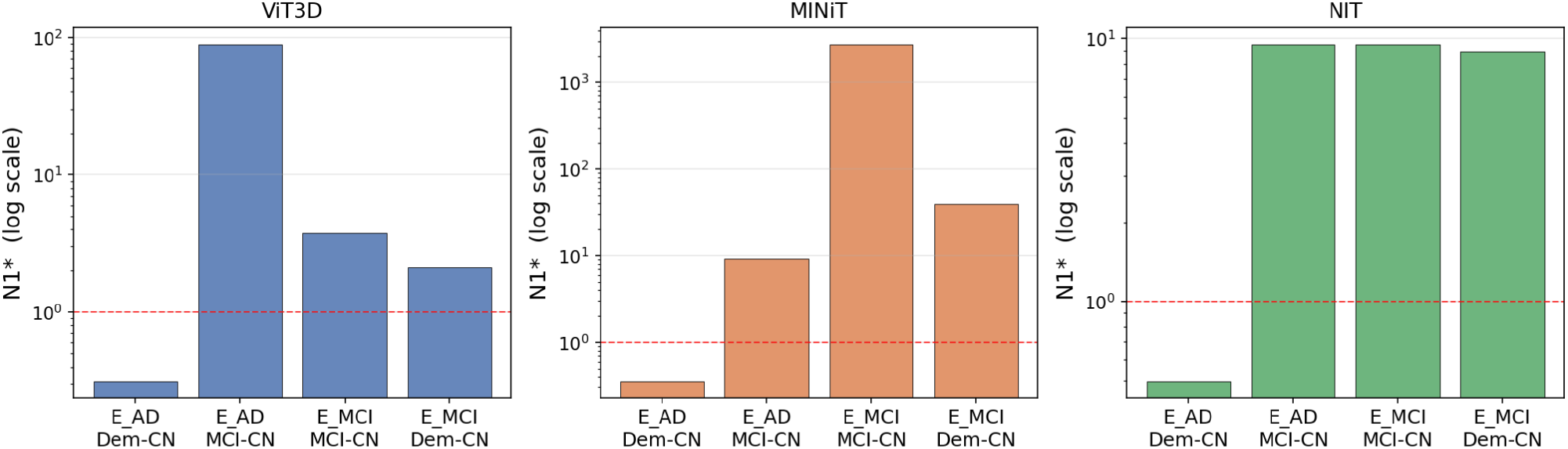
The 2×2 encoder × contrast design: N_1_* on a log axis, one panel per architecture (cross-fit, native basis; cells 2 and 4 leakage-controlled). For MINiT, cell 2 (frozen AD encoder, hard contrast, λ fixed) does not reproduce cell 3, while cell 4 (MCI encoder, easy contrast) stays well above cell 1 - the spectral geometry of the embedding depends on the encoder.

The absolute-scale columns show where the change is and is not. Retraining changes the overall scale of the embedding by a large factor: tr Σ and λ_1_ fall from ∼10–90 for the E_AD encoder (cells 1–2) to ∼0.04–0.4 for E_MCI (cells 3–4), a drop of roughly 200–2000×. This scale change is not what separates the cells, because N_1_* is unchanged if the whole embedding is rescaled. What moves instead is the signal projection: α_1_ falls from 6.71 (MINiT cell 1) to 0.014 (cell 3) while the PC1 concentration f_1_ stays high. This is the rank-1 collapse measured in the next section. Two qualifications bound the claim, and both concern geometry stability rather than any single threshold value. For ViT3D the frozen swap produces no clear cell-2/cell-3 separation, because its geometry does not reproducibly collapse on MCI/CN to begin with; EVR@1 and effective rank vary widely across seeds; so there is no stable structure for the swap to move. For NIT the cell-2 and cell-3 distributions overlap across seeds, again reflecting seed-unstable geometry on this contrast rather than a reproducible effect. The stable, seed-consistent separation is therefore specific to MINiT, the architecture whose representation is reproducibly collapsed and is reported as such rather than as a uniform law.

### 4.3 The reproducible signature: a detached rank-1 spike

This section evaluates the spectral collapse using log-log plots of the encoder eigenvalues (or strictly speaking, the eigenvalues of the data Gram matrix after mapping the data through the encoder into the embedding space). We fit a straight line to eigenvalues 2-50 on a log-log plot (skipping PC1), extend it back to position 1, and compare the real first eigenvalue to that prediction: the ratio S is the ‘spike.’ S ≫ 1 means PC1 towers above the trend (collapsed); S ≈ 1 means it fits in (healthy). Fitting the bulk slope (eigenvalue decay) over k = 2–50 (excluding PC1) and measuring the spike ratio 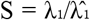 (bulk) shows that the three supervised Dementia-trained encoders carry a large, detached spike (S = 11.4–25.6), with subject-level bootstrap intervals far from 1, while the self-supervised DINO control shows S = 0.66, with bootstrapped confidence interval [0.59, 0.75] and excluding 1 from below (**Table 4, Figure 2**). In every supervised row 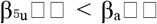 \, so PC1 pulled the overall slope upward and the steep supervised β is partly an artifact of fitting through the spike. Bulk R^2^ is 0.97–0.998, so directions 2–50 do follow a power law for the eigenvalue decay with respect to the eigenvalue rank, and the spike is a separate rank-1 feature. To better understand the same phenomenon, we can also compute EVR_1_ = 1/ζ_D(β), and this indicates the same effect: supervised encoders have a value that is 33–84% above their own power-law prediction, while DINO’s value is essentially still on the line (ratio 0.91). The spike is also a property of the encoder rather than the contrast. Routing the hard MCI contrast through the frozen AD encoder retains the large spike (cell 2: ViT3D 16.6, MINiT 10.3, NIT 8.5), and routing the easy contrast through the MCI encoder keeps its attenuated spike (cell 4: 4.0, 8.5, 4.6). Cross-fitting confirms this is a property of the covariance that is induced by the encoder, not an estimation artifact: the λ-side quantities that have to do with the covariance spectrum of the data Gram matrix (β, effective rank, EVR@1, λ_1_) are essentially invariant across cross-fit and closed-form estimators in every model×task cell, while only the discriminative signal side, and the values that depend on alpha, (i.e., α_1_, d□, N_1_*) vary. MCI-trained encoders show rank-1 (leading eigenvalue) spikes that are attenuated but still present (S = 3.4–9.4) rather than S ≈ 1; which is the no-spike case seen for the frozen encoder, DINO. A Ramsey RESET test [20], which refits the log–log bulk regression with powers of its fitted values added and tests whether they are jointly significant, i.e. whether the fit departs from a straight line, finds significant curvature in most bulk spectra. The bulk is therefore close to, but not exactly, a power law (log-linear), consistent with the high but imperfect fit quality (R^2^ = 0.97–0.998).

**Table 4.** Quantifying the rank-1 spike in the eigenvalue spectrum of the data Gram matrix, after mapping the data through different types of encoders. For each encoder, we fit the eigenvalue power law λ_i_ ∝ i^−β^ over the bulk of the spectrum (modes 2–50, excluding PC1) to obtain the bulk slope β_bulk, then measure the spike ratio 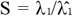 (bulk). This spike “size”, S, measures how many times larger the true PC1 is than the bulk fit predicted by extrapolation. A large S means that PC1 detaches from the rest of the spectrum (representation collapse); conversely, S ≈ 1 means PC1 is in line with the bulk (no collapse). We note two general observations: the **supervised encoders all have a prominent spike S** ≫ **1** (11–26 on Dem/CN) while the self-supervised DINO control does not (S = 0.66), and β_bulk < β_all in every supervised row, confirming that PC1 was the outlier steepening the naive slope. The final columns cross-check S against theory: EVR_1_ = 1/ζ_D(β) is the PC1 variance share a pure power law (where ζ is the Riemann zeta function), and the ratio of observed to predicted EVR_1_ indicates the same effect from the variance side. All quantities are computed on the training split; 95% CI from a 200× subject-level bootstrap; DINO is the negative control.

| Model | Task | $\beta_{\text{all}}$ | $\beta_{\text{bulk}}$ | Spike size, $S$ [95% CI] | $\text{EVR}@1$ obs | $1/\zeta(\beta)$ | ratio |
| --- | --- | --- | --- | --- | --- | --- | --- |
| VIT3D | Dem/CN | 2.312 | 2.063 | 25.6 [23.0, 27.1] | 0.955 | 0.701 | 1.36 |
| MINIT | Dem/CN | 2.271 | 2.052 | 17.4 [16.1, 18.6] | 0.941 | 0.691 | 1.36 |
| NIT | Dem/CN | 2.002 | 1.816 | 11.4 [10.3, 12.1] | 0.897 | 0.610 | 1.47 |
| VIT3D | MCI/CN | 1.755 | 1.660 | 3.4 [3.0, 3.7] | 0.762 | 0.517 | 1.48 |
| MINIT | MCI/CN | 2.127 | 1.955 | 9.4 [8.4, 10.2] | 0.938 | 0.650 | 1.44 |
| NIT | MCI/CN | 1.385 | 1.280 | 4.0 [3.3, 4.4] | 0.634 | 0.345 | 1.83 |
| DINO | Dem/CN | 1.148 | 1.180 | 0.66 [0.59, 0.75] | 0.189 | 0.208 | 0.91 |

**Figure 2.**
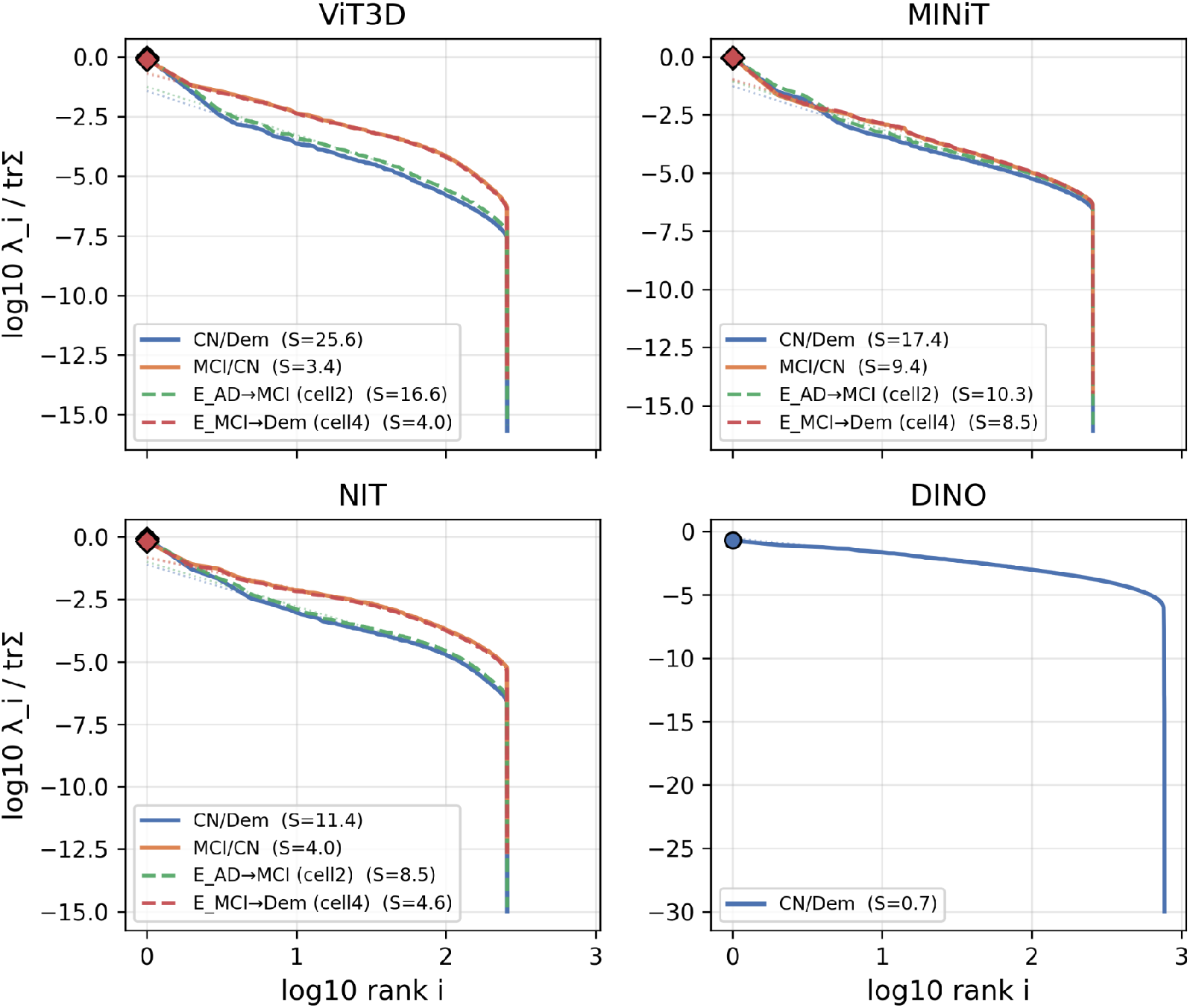
Eigenvalue spectra (log λ_i_/tr Σ vs log rank i) for each encoder, with both contrasts and the two frozen-transfer cells overlaid per panel. The dotted line is the power-law fit to the bulk of the spectrum (modes 2–50); the diamond marks PC1. The spike ratio S (legend) is how far PC1 rises above the bulk fit - S ≫ 1 means PC1 detaches from the rest of the spectrum (rank-1 collapse), S ≈ 1 means it lies on the bulk. The three supervised encoders show a detached PC1 spike (S = 11–26 on CN/Dem), and the spike travels with the trained encoder rather than the contrast: it persists when the frozen encoder is applied to the other contrast (E_AD→MCI, cell 2; E_MCI→Dem, cell 4). The self-supervised DINO control has no spike (S = 0.7) and a long, gently decaying spectrum instead. Tr Σ is normalised so panels are comparable across the ∼200–2000× scale differences between encoders.

### 4.4 The scaling curve rises with training sample size if the class-separation signal is distributed across the encoder eigenspectrum

The ‘forward model’ is the zeta law used as a predictive model for AUC(N): we input an encoder’s measured spectrum (λ_i_, α_i_) and it outputs a predicted accuracy curve. Here we test only its directional prediction that AUROC should rise with N when discriminative signal is spread across many modes, and stay flat when it is concentrated in PC1; not the exact shape of the curve, which the model overestimates (the red lines in **Figure 3** sit above the observed points, and the over-prediction is examined in next section). The directional prediction holds: for the one non-collapsed encoder, DINO [6] (β = 1.148, effective rank ≈ 52), observed AUROC rises from 0.683 at N = 50 to 0.793 at N = 2822, whereas the collapsed supervised encoders stay flat across the same range (**Figure 3**). We therefore validate the rising-vs-flat contrast, not a calibrated point-by-point fit. This reflects DINO’s distributed signal: d□ varies meaningfully with N rather than saturating at N = 50. For the supervised encoders, the large-sample prediction saturates to 1.0 at all observed N, a direct consequence of near-rank-1 collapse concentrating essentially all Mahalanobis signal in PC1 before N = 50 - the same collapse the spike ratio quantifies.

**Figure 3.**
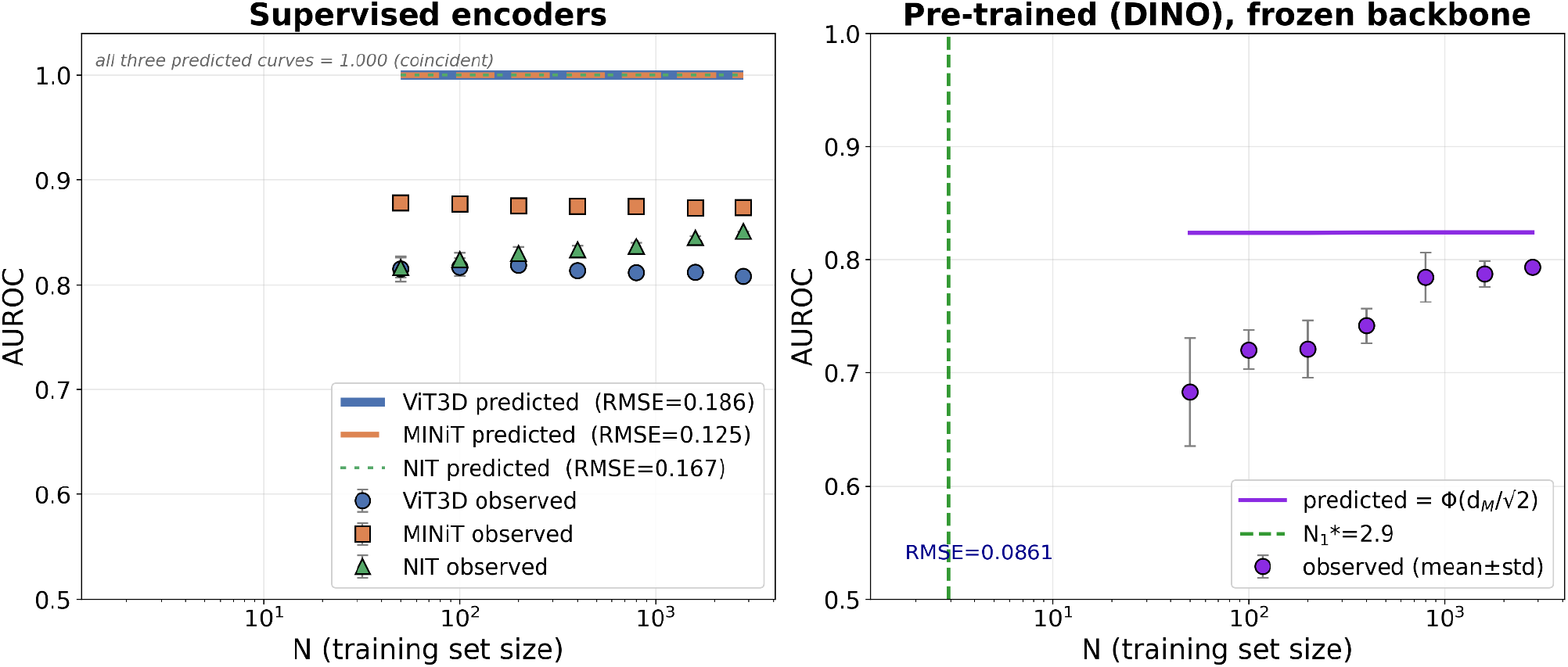
Testing whether AUROC rises with N. Observed AUROC vs training-set size (points) with the zeta law prediction Φ(d□/√2) (lines), for the three supervised encoders (left) and the frozen DINO encoder (right). The test is directional: when discriminative signal is spread across many modes, admitting more directions as N grows should make AUROC increase; when it is concentrated in PC1, AUROC should be flat. This holds as DINO’s observed AUROC rises from 0.68 (N = 50) to 0.79 (N = 2822), whereas the supervised encoders stay flat across the same range. We do not claim the prediction reproduces the curve: on the left it pins at its 1.0 ceiling (rank-1 collapse concentrates all signal in PC1 before N = 50, so there is no informative scaling to model), and for DINO it over-predicts at low N (purple line ≈ 0.82 while the observed points are lower), meeting the observations only at large N.

### 4.5. The apparent admission threshold is not an observable staircase

The zeta law for linear classifiers in *p* dimensions predicts that (1) AUC(N) will rise gradually with N, (2) there are staircase jumps when different eigenvalues of the data covariance can be stably estimated, (3) if the spectra for α and λ have been fitted, then we can estimate N_K*, which is an estimate of the amount of training data needed to stably estimate the k-th eigenvalue. In the current setting, we wanted to test if we could estimate N_K*, but in what follows we show that we could estimate it from the available data. For MINiT, we estimated N_1_ = 283.6, which may suggest we will see a staircase profile for AUC(N). A label-permutation null generates the distribution of N_1_ under “no signal, same covariance”; because N_1_ = λ_1_/α_1_^2^ has 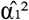 in the denominator, a near-zero true signal makes it very heavy-tailed. **Table 5** and **Figure 4** show the observed MINiT value lies inside the null band, below the null median, with empirical p = 0.94 (94% of label-shuffled runs give a larger N_1_); ViT3D and NIT fall below the entire null range (p = 1.0). Across 10 seeds MINiT’s N_1_ ranges 27.5–865.6 (split-half log_10_ SD = 0.78), so a single-seed value is one draw from a wide distribution and is not a stable estimand. The covariance geometry is, by contrast, reproducible for MINiT: EVR@1 stays 0.76–0.99 and effective rank near 1 across seeds - its representation is reliably near-collapsed, and only α_1_ (hence N_1_) wanders. For ViT3D and NIT both the threshold and the geometry are seed-dependent on this contrast (EVR@1 spanning 0.19–0.95 and 0.29–0.98).

**Table 5.** Identifiability of N_1_* (MCI/CN, training split). Permutation null: 500 label shuffles. Seed range: across 10 training seeds. Split-half log_10_ SD: 100 partitions.

| Model | Observed $N_1^*$ | Null median | Null 5th–95th pct | Emp. p | Seed range $N_1^*$ | Split-half $\log_{10}$ SD |
| --- | --- | --- | --- | --- | --- | --- |
| ViT3D | 3.6 | 2116.7 | 258.9 – 159,176 | 1.000 | 0.28 – 27.7 | 0.07 |
| MINiT | 283.6 | 2186.3 | 272.5 – 197,090 | 0.942 | 27.5 – 865.6 | 0.78 |
| NIT | 8.9 | 2013.8 | 242.1 – 365,904 | 1.000 | 1.2 – 28.6 | 0.11 |

**Figure 4.**
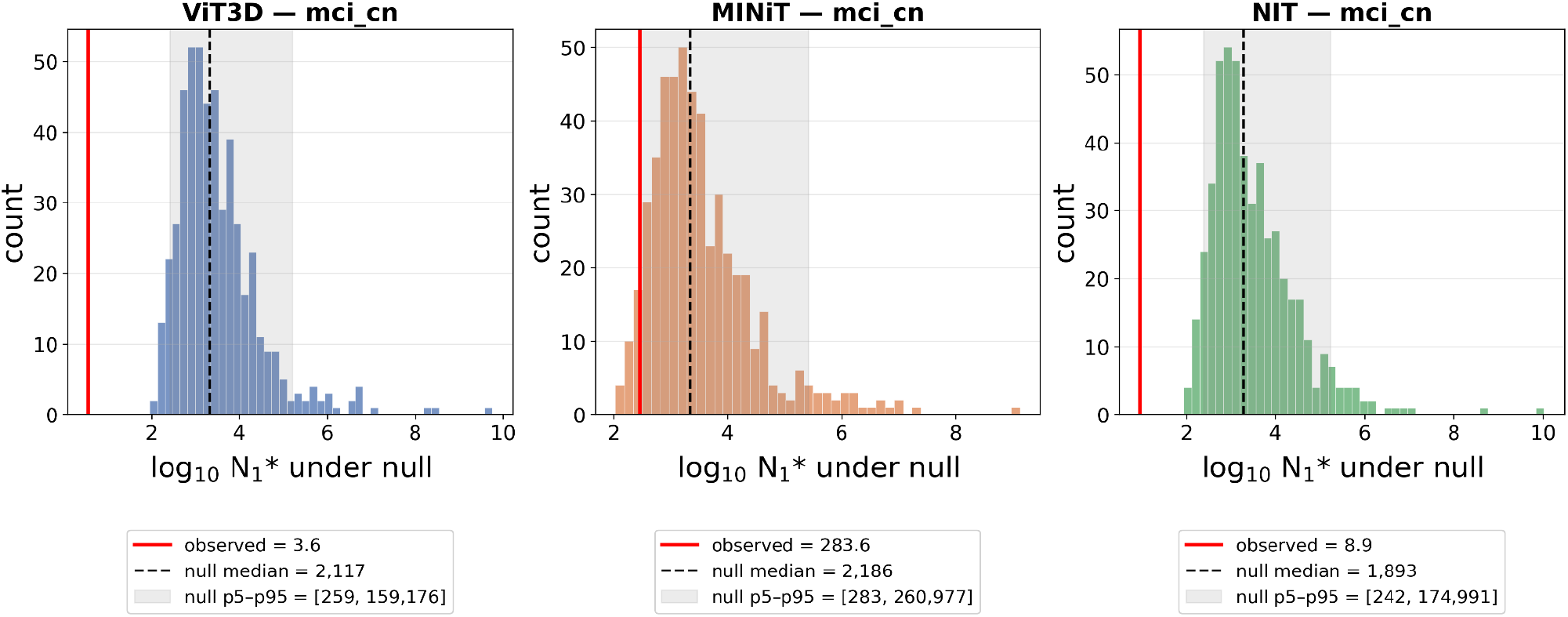
Permutation-null distribution of N_1_* under 500 label shuffles (MCI/CN, training split), one panel per architecture; the x-axis is log_10_ N_1_*. Each histogram is the N_1_* obtained when the diagnosis labels are shuffled i.e. the values expected under no class signal but the same covariance. The observed N_1_* (red) is compared against this null: for MINiT it sits below the null median (empirical p = 0.94, meaning 94% of shuffled runs give a larger N_1_*), and for ViT3D and NIT it falls below the entire null range (p = 1.0). In every case the observed N_1_* is not larger than what label-free noise produces, so on this hard contrast N_1_* is not distinguishable from a no-signal value. Legend values (observed, null median, null 5th–95th percentile) are raw N_1_*.

A matched empirical test agrees. The Zeta Law concerns a linear classifier, so we sweep N over {50–4057} with 10 subsampling seeds and a dense grid over 150–600, fitting a ridge linear probe (with LDA and 2-layer MLP checks) on the top-k(N) PCs and evaluating on the fixed test set (**Figure 5**). On the difficult MCI contrast, the probe does not perform better than chance (∼0.51–0.53) for all N as no direction ever carries an admissible signal and the MLP is equally flat, so this is not a limitation of the linear framing. The Zeta law prediction for AUC(N) over-shoots (0.69–0.93) and worsens as N grows, as k(N) climbs to ∼80 directions that carry no real signal. A formal staircase test (single-breakpoint vs smooth log model by BIC, with a bootstrap CI on the breakpoint) returns no step in any of the 12 cells: where a breakpoint improves the fit it captures gentle curvature, and its CI never contains the predicted N_1_*. The easy-contrast probes recover the end-to-end AUROCs (MINiT 0.873 by N ≈ 300, matching 0.878; ViT3D 0.808; NIT 0.848), confirming the pipeline. A natural worry is that this over-prediction is just the finite-sample inflation of Section A: because the class means are estimated from limited data, d□^2^ is biased upward, which would inflate the predicted AUROC on its own. But the inflation is far too small to explain the gap — the estimated bias in d□^2^ is 0.042–0.069, only 1–15% of the measured d□^2^, whereas the prediction overshoots the observed AUROC by 0.12–0.23. Removing the bias barely changes the fit (RMSE 0.176 → 0.169), so the over-prediction is a feature of the collapsed regime, rather than an estimation artifact.The reproducible object in this regime is the rank-1 spike as in section 4.3 and not an admission threshold.

**Figure 5.**
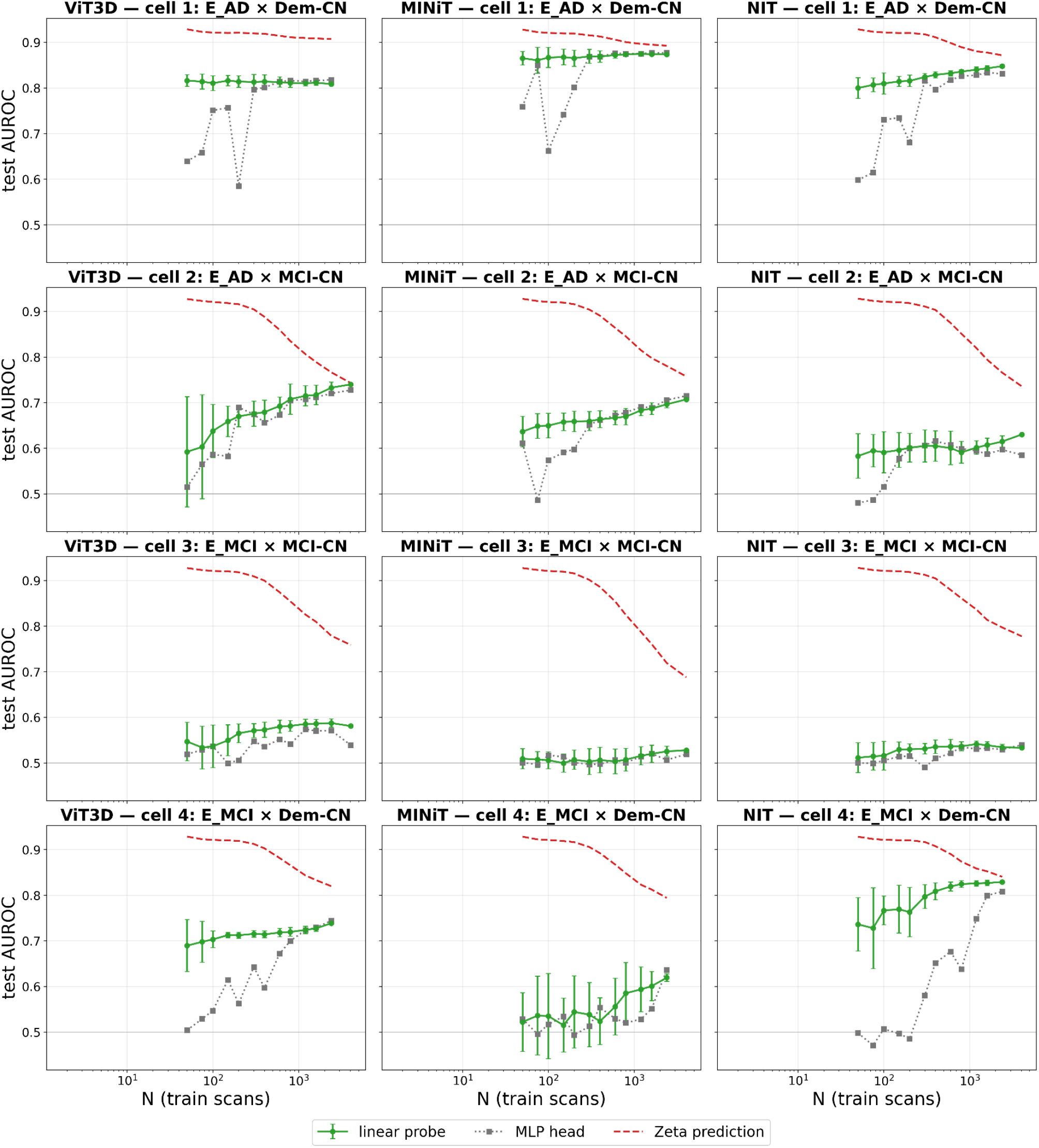
Frozen-encoder AUC(N): observed linear-probe (green) and MLP-head (grey dotted) curves with the Zeta Law prediction (red dashed), one panel per cell. The predicted curve is not monotone because the spectrum (λ_i_, α_i_) and the admitted set k*(N) are re-estimated on each size-N subsample, so the prediction carries its own finite-sample bias: the class-mean difference d is estimated from few scans at small N, so the plug-in Mahalanobis distance is inflated by a term scaling as 1/N (Section A). On the hard contrast, where the true signal is near zero, d□^2^ is almost entirely this inflation, pushing the predicted AUROC up toward ∼0.9; as N grows the inflation decays like 1/N, d□^2^ deflates toward its true near-zero value, and the prediction falls back toward the observed AUROC. So the decline in the red line is finite-sample inflation decaying with N — the per-N estimate of the model’s input is biased high at small N and corrects downward as N increases, not the model predicting that more data hurts. On the hard MCI contrasts (cells 2–3) the observed AUROC stays near chance at all N while the prediction starts far too high and declines toward it, meeting only at large N, and no breakpoint coincides with a predicted N_1_*. The linear probe tends to beat the MLP head at small N (fewer parameters to fit stably), with the MLP catching up as N grows.

### 4.6. Localizing the effect: the transformer, not the pooling

MINiT differs from the other architectures in how it produces the single embedding used for classification. ViT3D and NIT read out a dedicated CLS token, whereas MINiT aggregates its block tokens with ABMIL (attention-based multiple-instance learning), which is a learned, attention-weighted average of the per-block feature vectors. Given block embeddings h_1_, …, h_B (here B = 252), ABMIL computes a weight a_i_ for each block and returns the pooled embedding z = Σ_i_ a_i_ h_i_, where a_i_ = softmax(w^⊤^ tanh(V h_i_)) and V, w are learned parameters; the weights are non-negative and sum to one, so z is a convex combination of the blocks and the attention a_i_ says how much each block contributes. Because this pooling produces the embedding on which all our spectral quantities are computed, rather than sitting downstream of it, it is a candidate site for the collapse, so we ask whether retraining the pooling alone reproduces the effect.Three variants on MCI/CN (**Table 6**): V1, transformer and ABMIL both frozen from E_AD (the α-only condition); V2, transformer frozen, ABMIL retrained (17,281 of 3.27M params trainable); V3, everything retrained. In all 10 seeds V2 sits next to V1 and far below V3 (median N_1_ 1.74, 1.94, 65.5), recovering none of the V1→V3 change. As the pooling adapts to the new task, attention entropy rises 4.43 → 5.12 and top-10 block mass falls 0.440 → 0.252, most of the way from E_AD toward V3, yet N_1_ does not move (**Figure 6**). A linear probe on the frozen embeddings agrees (V1 0.708 ≈ V2 0.709 ≫ V3 0.564): retraining the pooling leaves linearly decodable signal essentially untouched, whereas retraining the transformer destroys most of it. The effect requires retraining the transformer feature extractor, where α_1_ collapses—consistent with the rank-1 collapse of section 4.3.

**Table 6.** MINiT ABMIL pooling ablation on MCI/CN (medians across 10 seeds). Linear-probe AUROC is subject-level grouped 5-fold on the training split; attention statistics over 252 blocks (uniform reference: entropy 5.53, top-10 mass 0.040).

| Variant | Configuration | $N_1^*$ | EVR@1 | erank | Probe AUROC | Attn entropy | Top-10 mass |
| --- | --- | --- | --- | --- | --- | --- | --- |
| V1 | Transformer + ABMIL frozen (E_AD) | 1.94 | 0.901 | 1.63 | 0.708 | 4.43 | 0.440 |
| V2 | Transformer frozen, ABMIL retrained | 1.74 | 0.850 | 1.81 | 0.709 | 5.12 | 0.252 |
| V3 | Fully retrained (E_MCI) | 65.5 | 0.983 | 1.12 | 0.564 | 5.42 | 0.070 |

**Figure 6.**
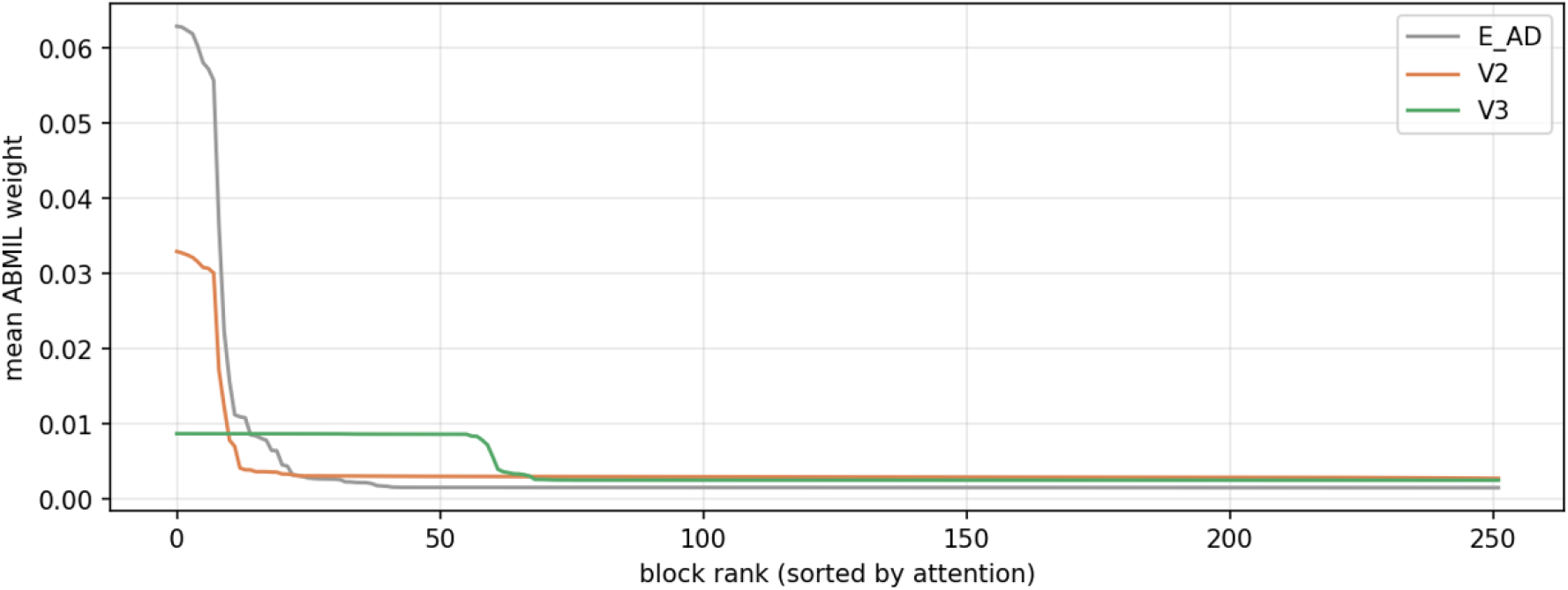
Retraining the pooling changes how it attends but not what the classifier can read out. MINiT aggregates its ∼252 block tokens into one embedding using ABMIL, a learned attention-weighted average; each curve shows how that attention weight is distributed across blocks (sorted, highest to lowest) for three conditions: V1 = both the transformer and its pooling frozen from the Dementia-trained encoder (E_AD); V2 = transformer frozen but pooling retrained on MCI; V3 = fully retrained on MCI. A steeper curve means attention is concentrated on a few blocks; a flatter curve means it is spread more evenly. The V2 curve moves substantially away from V1 (attention entropy 4.43 → 5.12, top-10 block mass 0.44 → 0.25), most of the way toward V3; so the pooling adapts to the new task. Surprisingly, this adaptation is not beneficial: V2’s N_1_* and linear-probe AUROC remain at their V1 (frozen) values, and only full retraining (V3) alters them (Table 6). The rank-1 collapse is therefore produced upstream, in the transformer that generates the tokens, not in the pooling that combines them.

## 5. DISCUSSION

### A. The scaling behavior, AUC(N), is a property of the learned representation

Overall, the scaling behavior of the different transformer encoders can be understood by examining the eigenvalue plots in **Figure 2**. First of all, when the data are mapped through any encoder, the data end up in an embedding space and the eigenvalues of its data matrix can be computed, and these indicate the effective dimensionality of the data after it has been fed through the encoder. The rate at which the eigenvalues decay depends on the encoder, and we were able to show that the supervised encoders (see the 3 left panels of the 4 in **Figure 2**) have a marked “spike” at the first eigenvalue, meaning that they have concentrated the covariance spectral energy to distinguish the classes in such a way that the encoder has a large first eigenvalue. In the case of the frozen encoder, DINO, the embedding is not tuned for the task and there is no rank-1 spike in the encoder eigenspectrum. This can be seen clearly in **Figure 2**, where the first (rank 1) eigenvalue stands higher than the linear trend that would be fitted through eigenvalues of rank 2 and higher.

All of the encoders, after training, have a spectral decay profile for their covariance geometry λ. Specifically after training, the data mapped through the encoders has a Gram matrix whose eigenvalues decay according to a power law - this can be seen from the fact that the eigenvalues decay roughly linearly on the log-log plots in **Figure 2**, except for the detached rank-1 spike, which is found in the supervised encoders, but not the frozen encoder (DIMO). This spike is produced by supervised training of the transformer on a fittable contrast: the amount that the first eigenvalue stands out is largest for the AD-trained encoders (S = 11–26), attenuated but present on the non-separable MCI contrast (S = 3.4–9.4), and absent in the self-supervised DINO control (S = 0.66). In linear discriminant analysis, this feature also arises and is called a “spiked covariance model” if the overall covariance of the cases and controls is used. The frozen-encoder design shows that the eigenvalue decay (spectral geometry) of the Gram matrix is a feature of the type of encoder (supervised or frozen), not the classification contrast: the spiking effect that appears when a model is retrained does not arise if we simply swap the contrast under a frozen encoder, nor by retraining the pooling alone. As such, the first major observation from our experiments is that supervised training concentrates the CLS representation into a rank-1 spike aligned with the between-class direction, and that the size of this spike scales with how separable the training contrast is: it is largest for the easily separable Dementia-vs-CN contrast (S = 11–26), attenuated for the harder, near-chance MCI-vs-CN contrast (S = 3.4–9.4), and absent for the self-supervised DINO encoder, which is not trained on a class contrast at all (S = 0.66).

### B. N_1_* is unreliable when the classifier is performing near chance

In the zeta law, **N**_1_**\*** is a measure of the sample size needed to stably fit the first eigenvalue of the data Gram covariance after embedding the data using the encoder. We found that this parameter can only be estimated reliably when the classifier performs better than chance, so it is not reliable when the classifier performs at near chance levels (i.e., when the training N is small or classes are hard to separate). The reason for this is that N_1_ = λ_1_/α_1_^2^ is intrinsically heavy-tailed when α_1_ is small, so it will not be stably estimand when the classifier is performing in the near-chance regime. If we do try to estimate this at low training N and with a difficult contrast, the single-seed values for **N**_1_**\*** cannot be interpreted as accurate admission thresholds. In future theoretical work, we will try to estimate the finite sample confidence intervals for the admission thresholds. This is an open mathematical question but may be tractable in the linear case, such as LDA in high dimensions. Two lines of evidence show that **N**_1_**\*** is unstable at low training N and for difficult tasks: the permutation null and seed variance, and the matched frozen linear-probe sweep, which do not detect any staircase step in the AUC(N) for any of 12 cells. All N_1_ statements are therefore reported as medians with dispersion. Conversely, the zeta law for AUC(N) predicts both a rate of increase with N and admission thresholds **N_k\*** only when the classifier is already performing better than chance.

### C. Qualifications

The large gap between cell 2 (the frozen Dementia-trained encoder applied to the harder MCI contrast) and cell 3 (the same architecture fully retrained on MCI) is specific to MINiT. ViT3D has no sizeable native MCI/CN threshold to begin with, and for NIT cell 2 and cell 3 are comparable; neither architecture settles to a stable spectral shape across seeds on the MCI contrast, so neither has a reproducible threshold for the frozen swap to separate. The separable/non-separable distinction is a matter of degree: the MCI-trained encoders retain a spike, attenuated to S = 3.4–9.4 rather than removed, and only DINO drops to S = 0.66. The bulk spectrum is well fit by a power law (R^2^ > 0.97) but not exactly log-linear, with a RESET test detecting significant curvature. The neural-collapse account is therefore strongest for the collapsed MINiT and AD-trained encoders and weaker, though still present, elsewhere.

### D. Limitations

Several limitations should be noted. First, the forward model systematically over-predicts AUROC in the near-chance regime: on the hard MCI contrast the predicted curve sits well above the observed one (**Figure 3, 5**). The main driver is finite-sample inflation of the Mahalanobis distance. Each direction admitted by k*(N) contributes α_i_^2^/λ_i_ to the predicted d□^2^, but when the true signal is near zero, the estimated α_i_^2^ is dominated by sampling noise rather than real signal. So as N grows and k*(N) admits ∼80 additional directions, each adds a small positive amount of fake distance and the prediction drifts upward toward its ceiling while the true AUROC stays at chance. Cross-fitting (Section C) rules out one version of this; bias from estimating d and the eigenbasis on the same data; but not the rest: the Gaussian/linear assumption in Φ(d□/√2) may not hold near chance, and repeat scans of the same subject inflate the effective N, so the admission rule Nα_i_^2^/λ_i_ > 1 over-counts independent evidence. The exact decomposition of these contributions is left to future work. Second, MCI vs CN AUROCs of 0.515–0.579 are near chance and no clinical utility is claimed—the task is used here to illustrate how transformers learn on easy tasks (dementia classification) versus hard tasks (MCI v control classification at low training N). Third, the DINO control was pre-trained on ImageNet; its AUC ceiling of 0.793 may be due to the domain gap between training on natural images (photographs) and medical images. A reasonable next step would be to test DINO with domain-matched self-supervised pre-training (on MRI data), which may do better so long as there is enough data to outperform the standard DINO vision encoder. Fourth, all experiments here were performed on a single dataset (ADNI) and two binary tasks; in future, we will test the generalization of these findings to other datasets and multi-class settings.

### E. Implications for Study Designs

When deciding if a deep learning task will benefit more from more data or a better encoder, it is worth estimating the geometry of the embedded data representation (i.e. the eigenspectrum of the data matrix in the embedding space of the encoder) on a small pilot sample. According to spectral theory, a valuable quantity to check is the rank-1 spike in the log-log spectral decay plot of the eigenvalues. As this generally follows a power law, the spike can be identified on a log-log plot, as we see in **Figure 2**. Measuring β_bulk, EVR@1, effective rank and the spike ratio S on a pilot sample takes minutes. A large spike (S ≫ 1) means the representation has collapsed onto a separable contrast and additional data will not help; the lack of a spike at the first eigenvalue (as we saw for DINO) means that the signal is distributed and collecting more data can pay off. This overall spectral analysis of performance scaling with training data provides a geometry-aware complement to Cohen’s-d power analysis that we classically use to estimate the effect size for a univariate 2-class classifier. With these spectral tools, we can begin to explore whether an (architecture, task) pair is in an active or saturated scaling regime as more scans are gradually collected.

A natural question is whether rank-1 collapse is inevitable or a limitation of the architecture and readout. It need not be inevitable. When a class is internally multimodal, the task’s intrinsic solution rank exceeds one even if the class-mean difference is a single direction: consider a binary “is this a 0” task over MNIST digits {0, 1, 2}, where the negative class contains two distinct sub-modes (1s and 2s). The Bayes-optimal boundary then requires more than one direction, but a transformer with a single pooled CLS embedding and a linear (or single-MLP) head can only express a rank-1 discriminant, so it collapses onto the dominant mode and cannot recover the second. Two routes could lift this ceiling. Architecturally, a transformer with a higher-rank readout such as multiple pooled projections rather than one CLS vector, could retain additional modes; we would expect the second mode to become admissible only at larger N, so a genuinely higher-rank encoder should show a rising rather than flat AUC(N) where the single-CLS model saturates. More broadly, objectives that learn a joint subspace of rank greater than one as in CLIP-style cross-modal training, where the target domain is itself high-dimensional, should resist collapse by construction, because the representation is optimized to align with a multi-dimensional target rather than a single binary axis.

## 6. CONCLUSION

We studied which properties of the deep learning model affect the data-scaling law for supervised 3D MRI vision transformers. Via several experiments, we found that performance scaling depends primarily on the learned representation. Specifically, it depends on the eigenvalue decay profile for the Gram matrix of the data, after it has been passed through the trained encoder, or the frozen encoder. We designed a frozen-encoder 2×2 experiment that held the embedding covariance λ fixed, and by separating the effect of encoder training from the difficulty of the task, we were able to show that the spectral geometry depends on the data encoder rather than the classification contrast. A sign of representational collapse is a detached rank-1 spike in the eigenvalue log-log plot, which is a property of the encoder (the prominence of the spike, S = 11–26 for supervised Dementia-trained encoders; S = 0.66 for the self-supervised DINO control). We also found that the admission threshold estimates N_1_* are not reliable when estimated at a low training N or for a difficult task where the classifier performs at chance level. One might read MINiT’s value of N_1_* = 283.6 as a real admission threshold - the sample size at which the leading direction first becomes usable, and so the first point where a supervised model’s accuracy should visibly step up with more data. But by permutation testing, we found that this value is not statistically distinguishable from noise: when the diagnosis labels are shuffled, 94% of runs produce an equal or larger N_1_*. It is also unstable, ranging from 27 to 866 across training seeds. And when we actually vary the training-set size, no step in accuracy appears. In this collapsed regime, then, the usable signal is better read from the shape of the eigenspectrum than from a single threshold value. The forward model runs the Zeta Law in prediction mode, from the measured spectrum (λ_i_, α_i_) it computes the admitted directions k*(N), the Mahalanobis distance d□, and a predicted curve AUC_pred(N) = Φ(d□/√2). Its directional prediction; AUROC rising with N when signal is distributed, flat when concentrated in PC1; was supported for DINO (observed AUROC rises with N) but not the collapsed supervised encoders (flat). We also showed this scaling happens due to training of the transformer and not due to the attention pooling architecture used by MINIT. Spectral collapse is thus a property of the learned representation, jointly determined by the training objective and by whether the training contrast is linearly fittable; a key practical recommendation for neuroimaging study design is to measure the rank-1 spike, at the current level of training data, to model whether additional training data will help. This is a real situation encountered by neuroimaging consortia and deep learning developers, as the added value of better encoders versus more training data is important to understand and estimate.

## Acknowledgments

This work was supported by NIH grants U01 AG068057, R01AG081571 and S10OD032285.

